# MAIT cells derived ligands signal via VEGFR2 to promote tissue repair and liver regeneration

**DOI:** 10.64898/2026.03.23.713159

**Authors:** Katia Sayaf, Martin Lett, Kate Powell, Irene Tasin, Lucy C. Garner, Aneesha Bandari, Narayan Rhamamurthy, Francesco P. Russo, Paul Klenerman, Carl-Philipp Hackstein

## Abstract

MAIT are a highly versatile population of innate-like T cells that have been implicated in promoting tissue repair-associated process in a variety of tissue and diseases settings in the last years. While certain specific effector molecules responsible for MAIT-cell mediated have been identified, the mechanisms by which MAIT cells exert repair functions remain incompletely understood.

Here, we show that hepatic MAIT cells express VEGFA, VEGFB and vimentin, an alternative ligand for the VEGFA-receptor VEGFR2 in both, regenerating and heathy tissue. Expression and secretion of these factors were induced *in vitro* by combined T cell receptor and cytokine stimulation. Supernatants of activated MAIT cells were able to promote proliferation of different epithelial and endothelial cells, including a liver sinusoidal endothelial-derived cell line in an VEGFR2-dependent manner. Together, our findings expand our understanding of MAIT cell function, especially in the liver and open new opens avenues for exploring MAIT therapeutic potential in modulating tissue repair.

## Introduction

Mucosal-associated invariant T (MAIT) cells are well-studied population of unconventional T cells which are defined by the expression of an invariant Va7.2-Ja33 (Va19-Ja33 in mice) invariant T cell receptor recognizing antigen including vitamin B2-derivates presented by the MHCIb molecule Major histocompatibility complex class I-related (MR1)^1–3^. MAIT cells can express a range of different effector molecules spanning classic Th1 and Th17 cytokines as well as mediators of cytotoxicity^4,5^ endowing MAIT cells with the ability to perform a variety of different functions in a disease- and tissue-specific context. Several studies have demonstrated that MAIT cells isolated from a variety of tissues including blood, skin and lung can promote processes associated with the repair of damaged tissues like proliferation, migration and differentiation of parenchymal cells both *in vitro* and *in vivo*^6–10^. Tissue repair is a complex phenomenon which is reflected by the genes included in different tissue repair signatures have been linked to the capacity of T cells to contribute to tissue repair over the years^11–13^ and contain a large variety of growth factors, proteases and other genes. These signatures have served as the basis to unravel the concrete pathways underlying the MAIT cells’ ability to promote repair as illustrated by the identification of Amphiregulin (AREG) as a MAIT-cell derived mediator of wound healing in the skin^9^ and tissue protection in autoimmune encephalitis^14^.

MAIT cells are massively enriched in the human liver^15^, and liver-derived MAIT cells have also been shown to upregulate repair-associated genes associated upon TCR stimulation *in vitro*^16^. As hepatic cells efficiently capture and present MAIT cell antigens from the circulation^17^, hepatic MAIT cells are in a highly stimulatory environment endowing them with the potential to contribute to the maintenance and regeneration of the liver. In line with this idea, the absence of MAIT cells aggravated pathology in a pre-clinical model of metabolic dysfunction-associated steatohepatitis (MASH)^18^, a finding that was linked to the MAIT cells’s ability to alter macrophage polarization towards an anti-inflammatory CD206+ M2 state. It is currently unknown however, which effector molecules are utilized by MAIT cells to exert tissue-protective and pro-regenerative functions in the liver.

Re-analysing publicly available data from regenerating human liver tissue^19^, we noted expression of different ligands for members of the vascular endothelial growth factor receptor (VEGFR) family by hepatic MAIT cells on the mRNA-level. VEGFR-ligands including VEGFA and VEGFB are part of the stablished tissue repair gene sigantures^11^ and upregulation of these growth factors on the mRNA level was observed in MAIT cells previous studies on the mRNA level^6,8,14^, however when attempted, so far, no expression was observed on the protein level^9^. VEGFA and VEGFB mainly signal via VEGFR2 and VEGFR1 homodimers respectivly^20^ and play different roles in tissue homeostasis with VEGFA-VEGFR2 interactions inducing angiogenesis, endothelial cell proliferation and migration, processes key for ensuring re-vascularization and blood supply of regenerating tissues. Signalling of VEGFB via VEGFR1 homodimers on the other hand exerts tissue protective functions by reducing oxidative stress^21^ and metabolic regulator^22^. The signalling network between VEGF and VEGFR family members is more complex though, as VEGFR1 as homodimer or heterodimer with VEGFR2 can also serve as a decoy receptor for VEGFA thereby finetuning VEGFAs pro-angiogenic function^20,23^, while it was also shown that Vimentin, a type III intermediate filament protein implicated in wound repair^24^, was shown to serve as an alternate ligand for VEGFR2, mimicking VEGFAs effects^25^.

Here, we demonstrate expression of the VEGFR-family ligands VEGFA, VEGFB and Vimentin in MAIT cells on the protein level. We further show that MAIT cells can impact tissue repair associated processes in an VEGFR2-dependent manner *in vitro* utilizing a variety of different assays. Our results expand the list of effector molecules produced by MAIT cells and demonstrate that they can directly promote the proliferation of different cell types, including liver sinusoidal endothelial cells (LSECs). These findings implicate MAIT cells as potential regulators of the re-vascularization of regenerating liver tissue.

## Results

### MAIT cells in the regenerating liver express VEGFR-ligands

To gain insights into possible MAIT-cell derived mediators that might promote repair-associated processes in the liver, we reanalysed an existing single-cell RNASeq (scRNASeq) derived from a patient cohort from a study made by Brazovskaja *et al.* that had undergone portal-vascular embolization^19^. In such patients, the blood flow to a disease part of the liver is blocked off several weeks prior to resective surgery, resulting in atrophy in the affected part (**Figure 1A**). In the remaining part of the organ however, the procedure induces a hypertrophic regenerative state resulting in the proliferation of the hepatocytes to compensate for the missing part of the tissue. To study the phenotype of MAIT cells in a tissue in which these repair-associated processes are taking place, we identified and analysed MAIT cells from the regenerating part of the liver and compared their transcriptional profiles to MAIT cells isolated from the atrophied embolised part or from healthy control tissues. Louvain-clustering revealed the existence of 4 separate MAIT clusters (**Figure 1B**). MAIT cells from the embolised tissue mainly populated cluster 0, while cells from healthy controls were contained in cluster 3 (**Figure 1C**). In contrast, MAIT cells from regenerating tissue were mainly localised in clusters 1 and 2 with some cells also scattered in the other clusters (**Figure 1C**).

**Figure 1.**
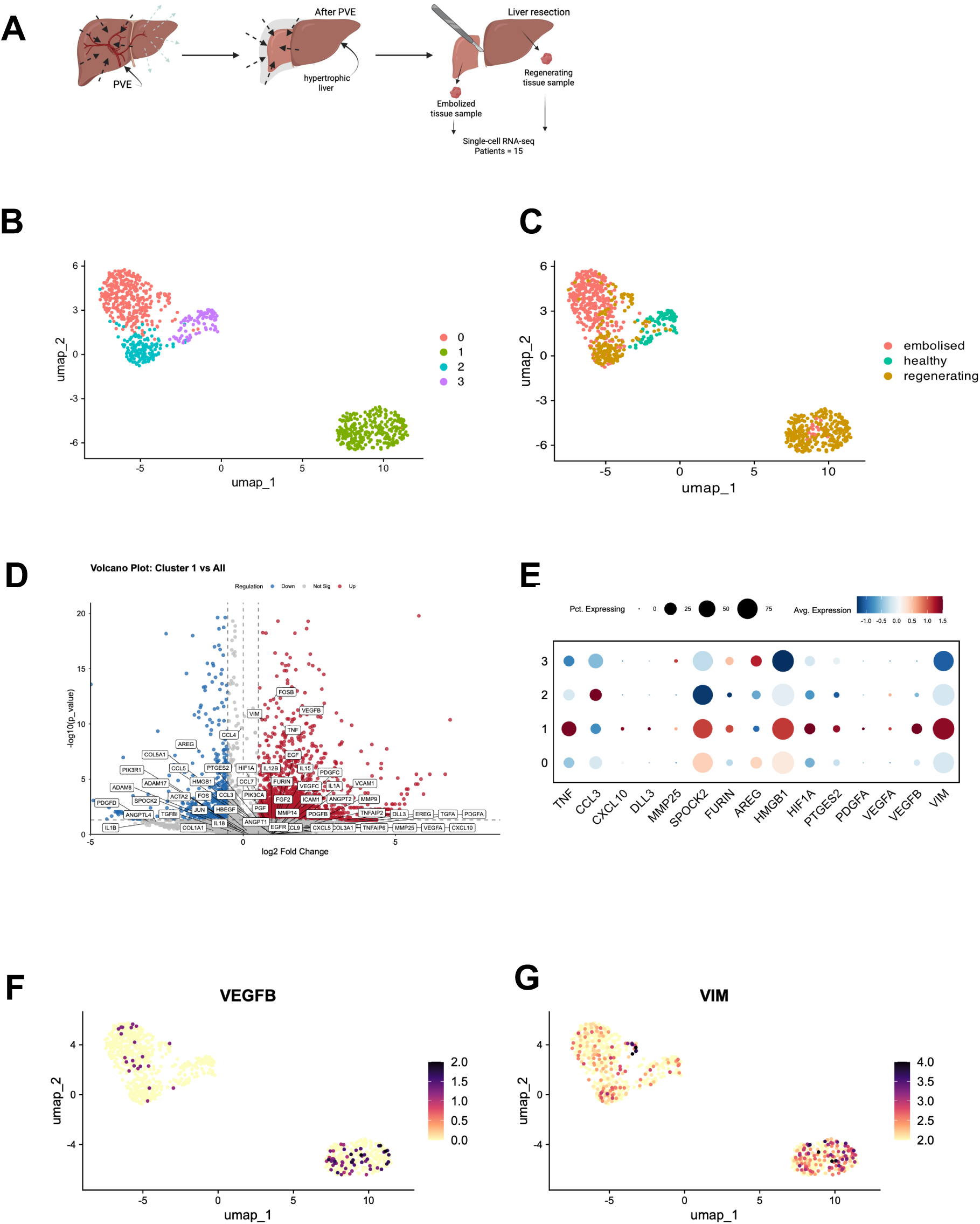
MAIT cells from regenerating human liver tissue show increased expression of VEGF and VIM. (A) Experimental design of the portal vein embolization (PVE) procedure as performed by Brazovskaja *et al*. (B) UMAP of Louvain clusters of hepatic MAIT cells from the indicated samples. (C) Origin of the cells in the clusters defined in B. (D) Volcano plot showing genes upregulated in Cluster 1 compared with all other MAIT clusters. (E) Dot plot displaying expression of selected tissue reapir-associated genes across the clusters defined in B. (F, G) Feature Plots showing the expression of *VEGFB* and *VIM* across MAIT cell clusters.

Gene expression analysis revealed the MAIT cells in cluster 1, which comprised exclusively cells derived from regenerating tissue, showed upregulation of several genes associated with tissue repair (**Figure 1D-F**) including HMGB1, HIF1A, FURIN and VEGFB. Interestingly, these cells also showed higher expression of VIM (**Figure 1E, G**), the gene encoding the intermediated filament protein vimentin. Interestingly, it has been demonstrated that endothelial and tumor cells can secrete vimentin upon which it can serve as an alternative ligand for VEGFR2, mimicking the proangiogenic effects of VEGFA^25^. Taken together, this analysis revealed that MAIT cells in regenerated liver tissue upregulate several mediators associated with repair-associated processes with a particular emphasis on ligands for members of the VEGF-receptor family like VEGFB and vimentin.

### MAIT cells require combined TCR- and cytokine signalling for the production of VEGFA, VEGFB and vimentin

To test whether MAIT cells have the capacity to produce VEGFR-ligands on the protein level, we next decided to model MAIT cell activation in vitro using blood-derived cells and MAIT cell lines. Our data show that upon stimulation the supernatant of sorted MAIT cells contained several different growth factors including VEGFA, PDGFA, GM-CSF and M-CSF (**Figure 2A**), while no differences were observed for several other growth factors (**Supplementary Figure 1A**). Among those, VEGFA expression in particular stood out as other T cells subsets sorted and stimulated in a similar manner failed to upregulate it to a similar extent (**Figure 2B, Supplementary Figure 1B**). This induction in general was seen at 48h – 72h post stimulation with a combination of TCR- and IL12+IL18 (**Figure 2A, B**). Expression of VEGFB was an exception to this as in contrast to VEGFA and other growth factors analysed we only observed VEGFB expression after a longer stimulation period of 6 days (**Figure 2C**).

**Figure 2.**
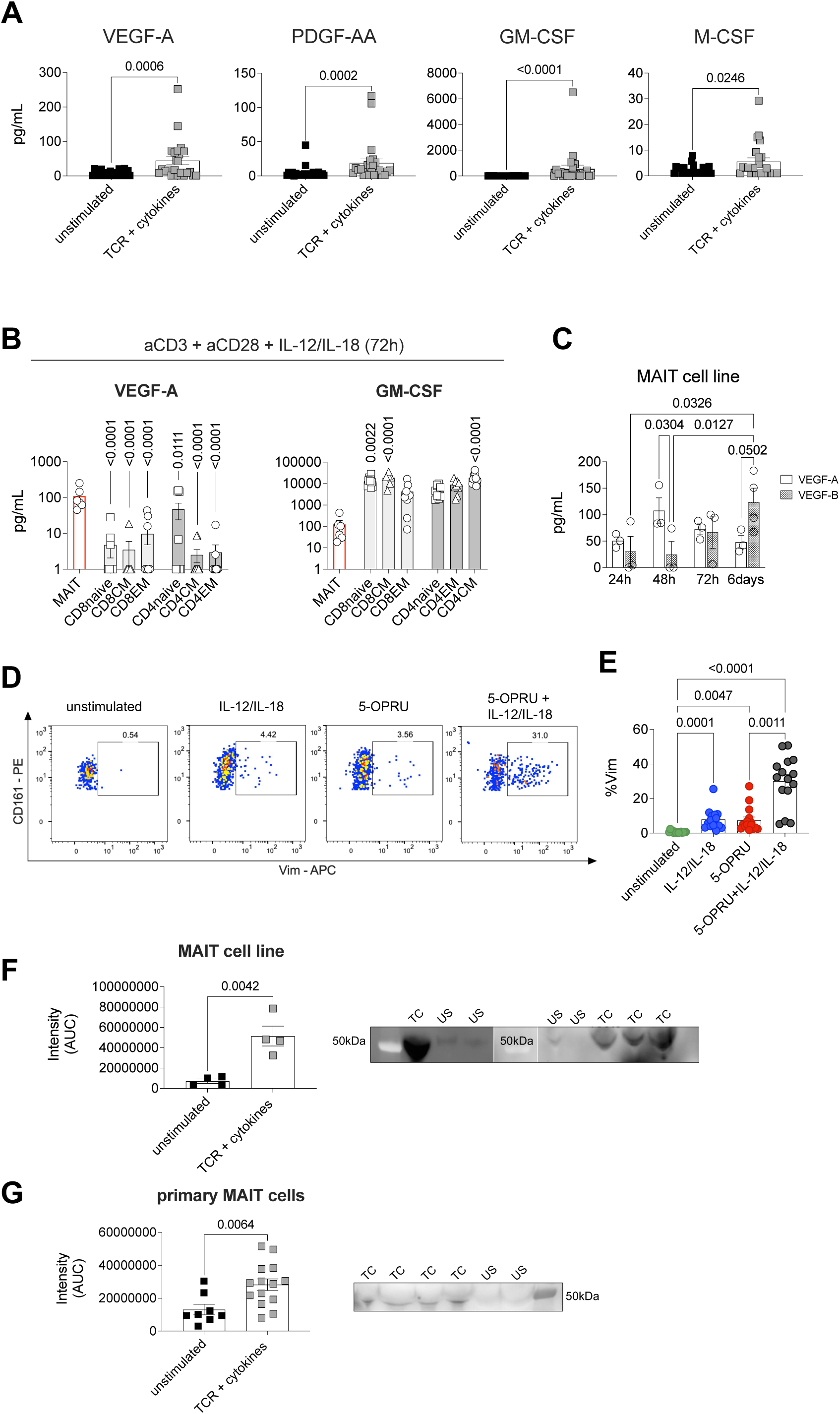
Activated MAIT cells secrete tissue repair–associated factors, including VEGF and vimentin. (A) Concentration of growth factors (VEGF-A, PDGF-AA, GM-CSF, and M-CSF) produced by primary blood-derived MAIT cells following activation with anti-CD3/anti-CD28 and IL-12/IL-18 for 72 hours, compared with unstimulated controls. (B) Concentration of VEGF-A and GM-CSF in the supernatants of activated MAIT cells or conventional CD8⁺ and CD4⁺ T cell subsets. Conventional T cells were classified as naïve (CCR7⁺ CD45RA⁺), central memory (CCR7⁺ CD45RA⁻), effector memory (CCR7⁻ CD45RA⁻), or terminally differentiated effector memory (CCR7⁻ CD45RA⁺). (C) Concentration of VEGF-A and VEGF-B from a MAIT cell line stimulated with PHA and IL-2 for 24, 48, and 72 hours, and 6 days. (D-E) Peripheral blood mononuclear cells (PBMCs) were left unstimulated or stimulated for 72 hours with IL-12 (50 ng/mL) and IL-18 (50 ng/mL), 5-OP-RU (10 nM), or a combination of both stimuli. Intracellular flow cytometry staining was then performed to assess vimentin expression in MAIT cells identified as live CD3+ CD161+ Va7.2+ cells. (F-G) Vimentin expression in primary MAIT cells isolated from PBMCs and in a MAIT cell line was assessed by Western blot and quantified using ImageJ. Cells were left unstimulated or stimulated with anti-CD3/anti-CD28 and IL-12/IL-18 for 72 hours.

To assess Vimentin expression in MAIT cells we first utilized flow cytometry. Parallelling our findings regarding the other factors, vimentin was strongly upregulated in MAITs after combined TCR- and IL12+IL18 stimulation (**Figure 2D, E**). We could further confirm the expression of Vimentin in MAIT cells using Image Stream analyses (**Supplementary Figure 1C**). Given that vimentin mainly functions as an intracellular filament protein and is upregulated upon cell activation intracellularly^26^ we next aimed to determine if vimentin was also secreted by MAIT cells. To this end we analysed the supernatants of stimulated primary MAIT cells (**Figure 2F)** or MAIT cell lines (**Figure 2G**) and detected Vimentin there as well, confirming its secretion. Secretion of vimentin in MAIT cells could be largely blocked with Brefeldin and monensin (**Supplementary Figure 1D),** while the blockade of an alternative secretion mechanism previously shown to be relevant in myeloid cells^25^ had only a small impact (**Supplementary Figure 1E**).

### MAIT cells in the human liver express VEGFA and vimentin in the steady state

So far, our data clearly demonstrate that blood-derived human MAIT cells have the capacity to produce different VEGFR-ligands upon stimulation *in vitro*. As it was reported by us and others^16,27^, tissue and circulating MAIT cells do differ transcriptionally, we next examined MAIT cells in the human liver. Importantly, as one of the aspects distinguishing liver and blood MAIT cells is the higher expression of activation-associated genes and pathways by the former, possibly as a result of the presentation of MAIT TCR ligands presented by different hepatic cell types^17^. Hence, we hypothesized that hepatic MAIT cells might express VEGFR-ligands already in the steady state. To test this, we assessed the expression of VEGFA and vimentin in hepatic MAIT cells via immunofluorescent imaging in normal human liver samples. We defined MAIT cells as CD3+ T cells co-expressing CD161 and TCR Va7.2 (**Figure 3A, B**) and assessed expression of Vimentin and VEGFA. Comparing individual MAIT cells across different images we found different expression combinations with some cells expressing both molecules, either or none of them (**Figure 3B**). Overall, as summarized in **Figure 3C**, the majority of hepatic MAIT cells were positive for both Vimentin and VEGFA, while subpopulations expressed only one of these molecules and a small fraction being negative for them. In sum, our data demonstrate that MAIT cells have the capacity to produce and secrete VEGFA, VEGFB and vimentin *in vitro* as well as in the human liver *in vivo*.

**Figure 3.**
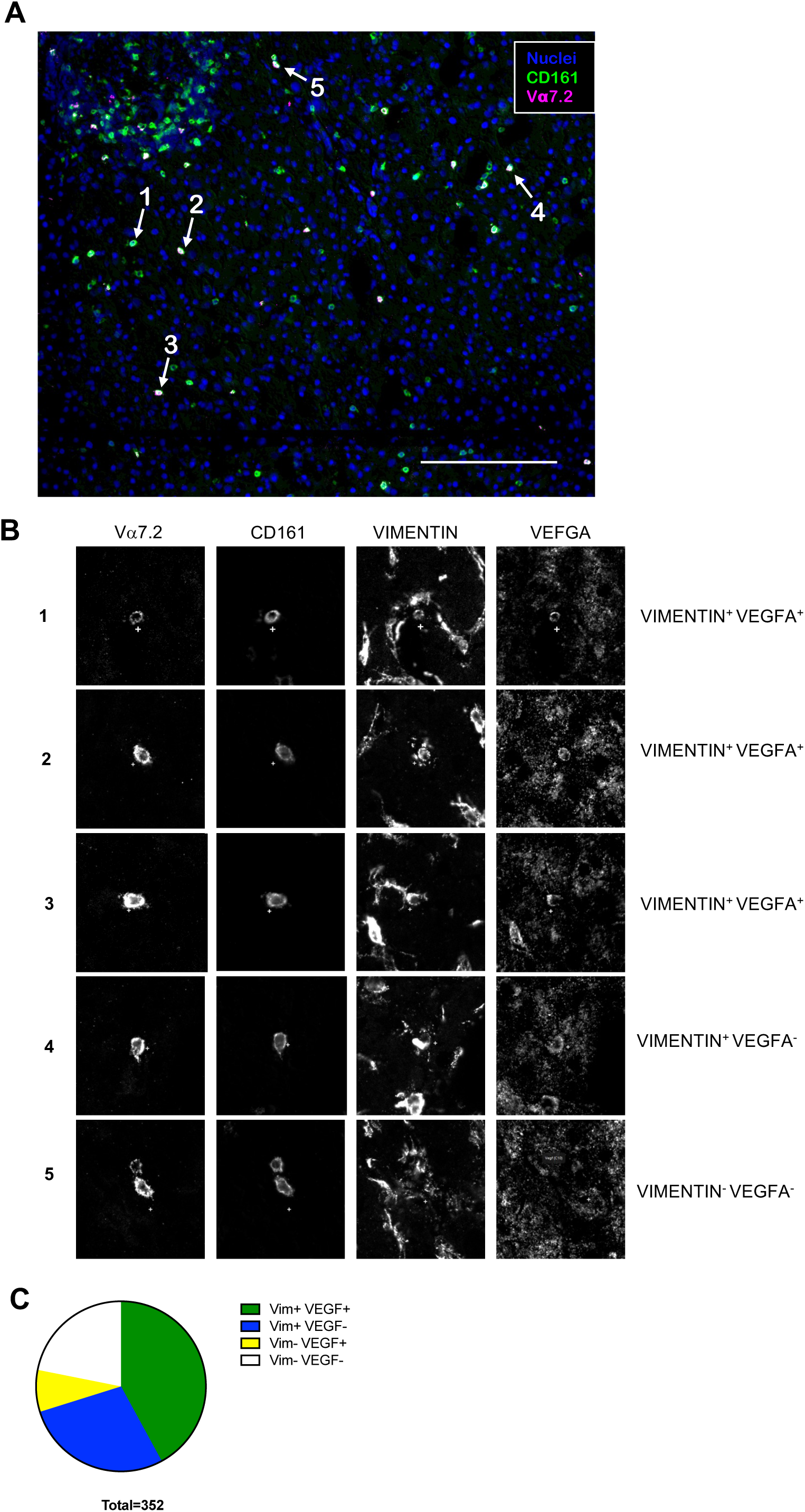
MAIT cells in the human liver express VEGFA and vimentin at steady state. (A) Representative images showing nuclear staining and expression of Vα7.2 and CD161 in fixed healthy human liver samples. Samples were mounted on cytometer chips and iteratively stained as described in the Methods section. Arrows indicate single cells analyzed below. (B) Single-channel images of vimentin, VEGFA, CD161, and Vα7.2 highlighting positive and negative fluorescence signals in MAIT cells within the hepatic environment as indicated in A. (C) Quantification of MAIT cell subsets based on vimentin and VEGFA expression. As shown in the graph, the majority of MAIT cells in the analysed liver sections are Vim⁺VEGFA⁺, with a substantial proportion of Vim⁺VEGFA⁻ cells. Scale bars, 50 μm.

### MAIT supernatants promote tissue repair-associated processes in a VEGFR2-dependent manner

We could previously show that the supernatants of MAIT cells can promote the migration and proliferation of Caco2-cells in *in vitro* wound healing and zone exclusion assays^6^. Both VEGFA and vimentin could increase area recovery in such assays we supplied as recombinant proteins in as dose-dependent manner and importantly, this effect could be abrogated by the addition of an antibody blocking VEGFR2 (**Supplementary Figure 2A-C**). To demonstrate the functionality of MAIT-derived VEGFR-ligands we next tested their capacity to promote repair-associated processes using *in vitro* zone exclusion assays in Caco2 as well as hepatic HHL12-cells. For this purpose, we collected the supernatants of FACS-sorted MAIT cells stimulated with the MAIT TCR ligand 5OPRU and IL12+IL18 for 72h or unstimulated controls. We then exposed the cell lines to these supernatants or culture medium and – based on the cell line – assessed area recovery at different time points (**Figure 4, Supplementary Figure 2D, E**). In both cell lines, the addition of the supernatants of stimulated MAIT cells significantly increased the rate of which the cells recovered the area they were previously excluded from (**Figure 4B, C, Supplementary Figure 2E**). This increase in recovery rate could be completely prevented by the addition of blocking antibodies targeting VEGFR2, the receptor for VEGFA and vimentin (**Figure 4B, C, Supplementary Figure 2E**). These data show that MAIT cells can promote the proliferation and migration of epithelial-type cells including hepatocyte-like cells in the VEGFR2-dependent manner.

**Figure 4.**
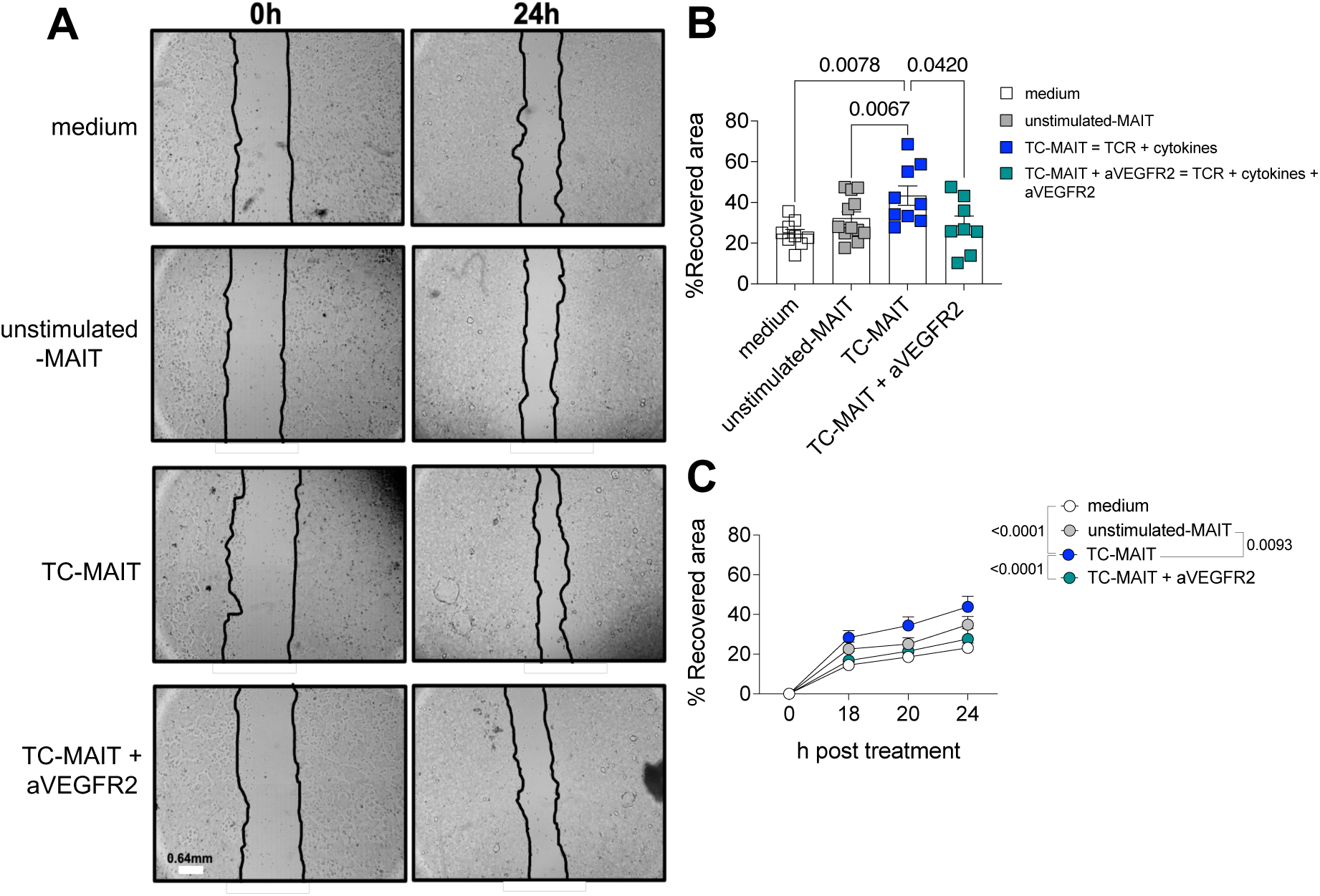
Activated MAIT cells promote wound healing *in vitro* in a VEGFR-2 manner. Primary MAIT cells were sorted from PBMCs and left unstimulated or stimulated with anti-CD3/anti-CD28 and IL-12/IL-18 for 72 hours, after which supernatants were collected and stored at −80 °C. Caco-2 cells were seeded in 24-well plates with culture inserts, and upon formation of a confluent monolayer, inserts were removed to generate a wound. Supernatants from unstimulated MAIT cells, stimulated MAIT cells, or stimulated MAIT cells in the presence of a VEGFR-2–blocking antibody were added. Images were acquired at 0, 18, 20, and 24 hours to assess wound closure. (A) Representative images of Caco-2 cell monolayers before (0 h) and following treatment with MAIT cell–derived supernatants with and without anti-VEGFR2 antibody. Magnification is 10X. (B,C) Quantification of wound closure, expressed as the percentage of recovered area, after 24 hours (B) and over the indicated time course (C) for each experimental condition.

Given that we could show that MAIT cells can produce both VEGFA and vimentin (**Figure 2, Supplementary Figure 1**) we next wanted to determine if the effects we observed are dependent on one of them or whether they work synergistically. To this end, we performed another set of *in vitro* zone exclusion assays testing the impact of a neutralizing anti-vimentin^25^ antibody by itself or in combination with anti-VEGFR2 on area recovery (**Figure 5, A-C**). The quantification of the results revealed that the neutralization of vimentin did significantly reduce area recovery, while the addition of anti-VEGFR2 did not have a significant additional effect (**Figure 5B, C**), indicating that Vimentin is responsible for the bulk of VEGFR2-dependent effects of MAIT cell supernatants on cell lines. Taken together these data suggest that production of vimentin by itself is sufficient to mediate the VEGFR2-dependent effects observed in our experiments.

**Figure 5.**
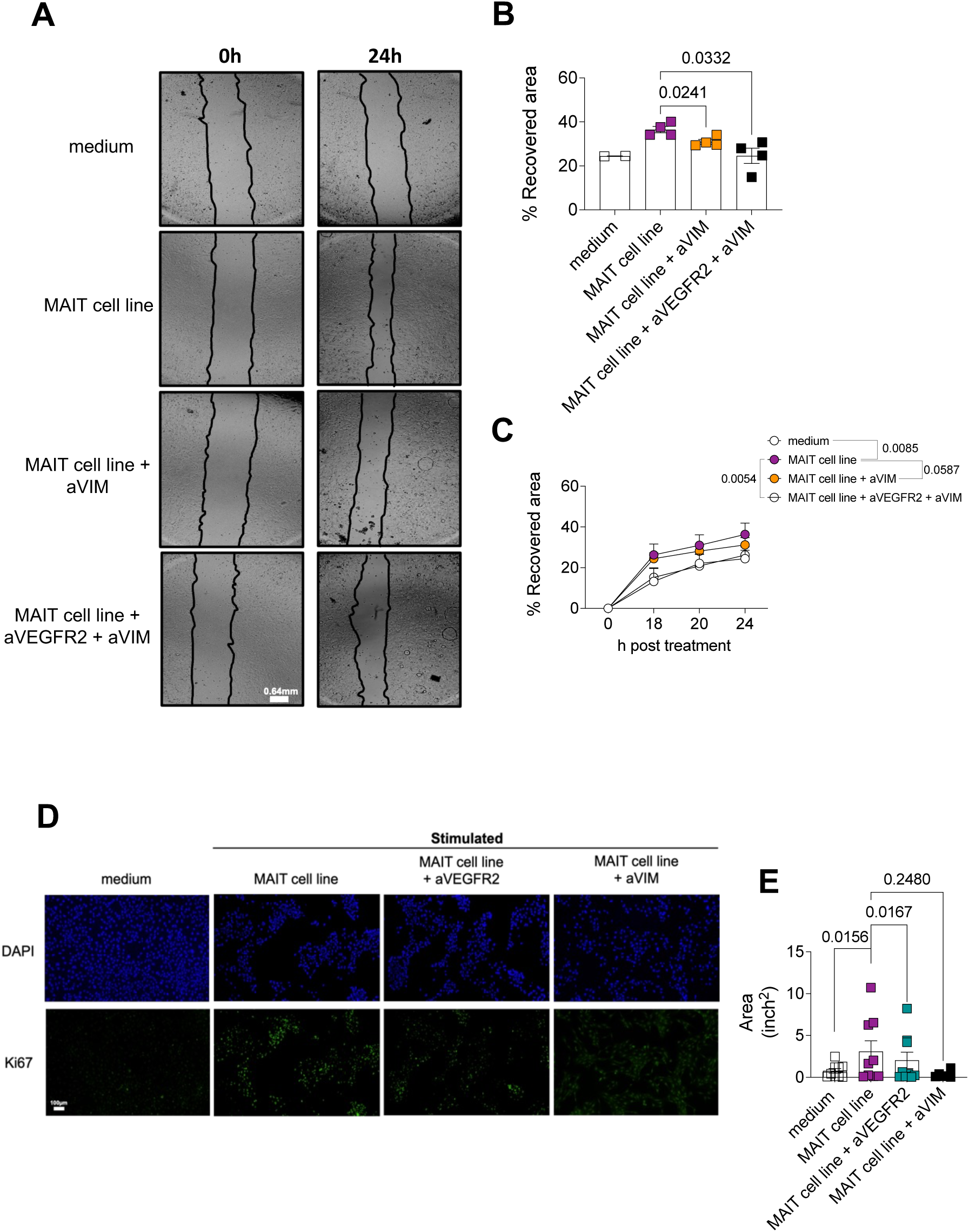
Vimentin production is sufficient to mediate the VEGFR-2–dependent effects of MAIT cells. (A) Representative images of Caco-2 cell monolayers at the time of scratch (0 h) and following treatment with MAIT cell–derived supernatants in the presence or absence of a VEGFR-2-blocking antibody. Magnification is 10X. (B,C) Quantification of wound closure, expressed as the percentage of recovered area, after 24 hours (B) and across the indicated time course (C) for each experimental condition. Data are representative of two independent experiments using cells from four donors each; each plate included two control wells. (D) Representative immunofluorescence images of Ki67⁺ liver sinusoidal endothelial cells (LSECs; TMNK1 cell line) cultured with or without MAIT cell line-derived supernatants, in the presence or absence of VEGFR-2- or vimentin-blocking antibodies. Images were acquired 24 hours after supernatant addition. Magnification is 10X. (E) Quantification of Ki67⁺ area in LSECs under the indicated conditions. Scale bar is 100μm.

### MAIT-mediated VEGFR2-signalling induces proliferations of LSECs

The key outcome of VEGFR2 is the promotion of angiogenesis via the activation of endothelial cells. This activation leads to the degradation remodelling of the surrounding extra-cellular matrix via endothelial cell-derived protease and the subsequent proliferation of endothelial cells. Hence, we also assessed the impact of MAIT-supernatants and VEGFR2-ligands in them on a LSEC - derived cell line (TMNK1), as LSECS are a major endothelial cell population of the liver.

Consistent with the expression of VEGFR2-ligands by MAITs, the supernatants of MAIT cell cultures were able to increase expression of the proliferation marker Ki-67 inTMNK1 cells (**Figure 5D, E).** Conversely, blockade of VEGFR2 or neutralization of vimentin prevented or dampened this effect (**Figure 5, E**). These data suggest that MAIT cells can induce the proliferation of LSECs.

## Discussion

The term tissue repair encompasses several complex processes that regulate the migration, proliferation and differentiation of parenchymal cells. Mirroring this complexity, gene signatures often used to demonstrate the ability of a cell population to contribute to repair contain a large variety of different proteins, including, growth factors, chemokines and cytokines, transcription factors and proteases impacting on a substantial number of different signalling pathways^11–13,28^. Here, we uncovered upregulation of VEGFA, VEGFB and vimentin in hepatic MAIT cells and PBMC-derived MAIT cells upon activation. While it is well recognized that VEGFA and VEGFB can contribute to repair processes, vimentin is best known for its role as a structural protein. However, vimentin also has been recognized as a repair-associated gene before^28^ and given its ability to mimic the effects of VEGFA-signalling^25^, we decided to examine the ability of MAIT cells to produce and secrete vimentin, as well as its impact on *in vitro* correlates of tissue repair and endothelial cell proliferation. A previous report demonstrated that while vimentin functionally mimics VEGFA, compared to the latter much higher concentrations of vimentin are required to induce sprouting of endothelial cells *in vitro*^25^. However, in the presence of recombinant vimentin, several VEGFA-induced effects including lower VE-cadherin-expression as well as higher VEGFR2-expression and phosphorylation were observed,^25^ suggesting that both of these factors work synergistically and vimentin might an important regulator of the VEGFA-VEGFR2 signalling axis. Secreted vimentin has also been reported to several other receptors including insulin-like growth factor 1 receptor (IGF1R)^29^,CD44^30^ and possibly Ryk^31^ further broadening the spectrum of possible target cells and effects of MAIT cell derived signals. Neutralization of vimentin reduced the effect of MAIT cell–derived supernatants to a similar extent as blockade of VEGFR2. This observation suggests that vimentin may account for a substantial fraction of the VEGFR2-dependent effects induced by MAIT cells, at least under the *in vitro* conditions used in this study. However, the existence of additional vimentin receptors beyond VEGFR2 raises the possibility that it promotes epithelial and endothelial cell proliferation through multiple signalling pathways. Such parallel signalling could partially mask effects mediated by VEGFA in our assays. Accordingly, our data do not exclude an important functional contribution of MAIT cell–derived VEGFA.

Our analyses also showed upregulation of VEGFB in the regenerating liver and, given enough time, upon stimulation in MAIT cells in culture. VEGFB signals via VEGFR1 rather than VEGFR2 and does not promote angiogenesis. However, in addition to reducing oxidative stress^21^, it has been described that VEGFB can induce the expression of hepatic growth factor (HGF) by LSECs or their precursors^32,33^. As HGF is an important mitogen for hepatocytes MAIT cells might indirectly promote hepatocyte proliferation.

PBMC-derived MAIT cells required strong stimulation to produce VEGFR-ligands, in line with the previously published data that indicate the MAIT cell tissue repair function are unlocked by TCR-stimulation, particularly in combination with strong cytokine-mediated signals^6,8,10,34^. Interestingly, when analysing human liver tissue, we were able to detect both VEGFA and vimentin in a substantial fraction of MAIT cells, suggesting that hepatic MAIT cells exists in an activated state. This is in line with previous findings showing the presentation of gut-derived MAIT-ligands by hepatic cells^17^ as well as transcriptional data showing upregulation of activation-associated genes and gene signatures in the liver compared to the blood^27^. Our data support the model that hepatic MAIT cells are continuously activated in the healthy tissue and contribute to tissue integrity^35^. This has important implications for the understanding of the pathomechanisms of chronic liver diseases, as depletion and dysfunction of MAIT cells has been noted in HBV^36^, HCV^37^ and other diseases associated with liver fibrosis including autoimmune hepatitis^38^. On the other hand, it is also known that MAIT cell activation in the liver can be detrimental in specific contexts as it observed that MAIT cells can promote liver fibrosis^38,39^ and could contribute to immune pathology during viral hepatitis^40^. It should be noted that MAIT activation in these disease settings appears to be primarily cytokine-driven^38^, and hence, lacks the TCR component crucial for many of the MAIT cells repair-associated effector functions^6,10,16^. Hence, a key objective of future studies would be to unravel the exact signalling mechanisms, i.e. the impact of different cytokines or TCR signalling strength and duration, that regulate MAIT cell polarization towards a tissue protective or a pro-inflammatory phenotype.

The ability of MAIT cells to promote tissue repair has been demonstrated *in vitro*^6^, in the context of infection^8^ and specifically during wounding of the murine skin^7^. Yet, with the notable exception of AREG^9^, the effector molecules used by MAIT cells to mediate tissue repair remain largely unknown. Our demonstrates that ligands of the VEGFR family are part of the MAIT effector repertoire. Importantly, given the complexity of the process of tissue regeneration further amplified by tissue-, disease and timepoint-specific aspects, we predict that MAIT cells do not produce all potential repair associated factors in their repertoire at the same time. As it was shown for other MAIT effector molecules including members of the IL-17 family^27^ and also demonstrated here for MAIT-derived VEGFA and VEGFB, there are clear temporal patterns: MAIT cells produce VEGFA relatively early and in a transient manner after activation, while upregulation of VEGFB expression require prolonged periods of activation and - at least *in vitro*- can only be observed after 6 days. From a biological point of view these different phases suggest a model in which MAIT cells early on after activation produce factors that can promote the proliferation of target cells like VEGFA. Prolonged activation on the other hand induces the production of factors like VEGFB which is implicated in promoting the survival of already differentiated cells, e.g. by reducing oxidative stress or inducing other factors like HGF to promote tissue regeneration. Hence, during liver injury, MAIT cells initially could promote the revascularization of the regenerating liver tissue and later contributing to stabilize and maintain the freshly differentiated endothelial cells, while also providing indirect mitogenic stimuli to hepatocytes. Overall, our study expands the spectrum of known MAIT cell effector molecules and specifically connects them to processes relevant for liver regeneration and angiogenesis.

## Materials and methods

### Peripheral blood mononuclear cell (PBMC) isolation

PBMCs were isolated from fresh whole blood from healthy donors by layering Lymphoprep on top of the blood and density gradient centrifugation (Lymphoprep, Axis-Shield) at 931 g for 30 min with no brake. After centrifugation, the PBMC layer was collected and washed with R10 medium [RPMI 1640 (Sigma-Aldrich) enriched with 10% fetal bovine serum (FBS; Sigma-Aldrich) and 1% penicillin–streptomycin (PS, Thermo Fisher Scientific)]. Cells were washed twice with R10 medium before cryopreservation in freezing medium [90% FBS and 10% dimethyl sulfoxide (DMSO; Sigma-Aldrich)] and stored in liquid nitrogen.

### Sorting and stimulation of MAIT and conventional T cells

Before sorting MAIT cells from PBMCs, enrichment of CD8+ was performed by using a CD8 positive selection isolation kit (Miltenyi). CD8+ enriched cells were stained with CD8-PECy7, CD161-APC, CCR6-BV421, nIR dead dye for 30 minutes at 4 °C, washed twice and resuspend in MACS Buffer [PBS supplemented with 1% FBS and 1mM EDTA (Thermo Fisher Scientific)] for sorting. MAIT cells, identified as CD161+ CCR6+, were sorted by using a MA900 Cell Sorter in a sorting buffer [(PBS Ca++/Mg++ free, 25mM HEPES, 10% FBS(Thermo Fisher Scientific)]. Conventional T cells were sorted from whole PBMCs stained with CD4-APC, CD8-PE/Vio770, CCR7-FITC, CD45RA-PE/Dazzle594 and nIR live/dead dye and otherwise treated the same as the MAIT cells.

After the sort, all cells were washed and resuspended in R10. Purity check was performed after sorting and surface stained with CD3-PerCP-Cy5.5, CD161-PE, Va7.2-PECy7 (BioLegend) for MAIT or CD4-APC, CD8-PE/Vio770, CCR7-FITC, CD45RA-PE/Dazzle594 for conventional T cells. Samples were run at the MACSQuant (Miltenyi), revealing a purity of 94-98% for MAIT and between 90-96% for the conventional T cells.

### *In vitro* activation assays

For the assay 2.5×10^5^ sorted CD8+ primary MAIT cells or from the MAIT cell line were seeded on a Maxisorp 96-well-F-bottom plate (Thermo Fisher Scientific) coated with aCD3 (1.25µg/ml; BioLegend), and IL-12 (50ng/mL; Miltenyi) and IL-18 (50 ng/mL; MBL) together with aCD28 (1ug/mL; BioLegend), or PBS-treated wells were used as unstimulated controls. Supernatants were collected after 72h and stored at −80°C for further testing.

For the MAIT cell line activation assay, 2.5 × 10⁵ cells were seeded and stimulated with PHA and IL-2 for 24, 48, and 72 hours, as well as 6 days. Supernatants were then collected at each time point and stored at −80 °C for subsequent analyses.

For PBMC activation, 1×10^6^ cells stimulated for 72 h with IL-12 (50 ng/mL; Miltenyi) and IL-18 (50 ng/mL; MBL), 10 nM 5-OP-RU (kindly provided by David Fairlie), or a combination of 5-OP-RU and cytokines to achieve TCR- and cytokine-mediated activation.

### Flow cytometry

Brefeldin A (eBioscience, 1000×) was added together with Monensin (BioLegend, 1000×) where indicated; otherwise, Brefeldin A alone was added for the final 4 h prior to intracellular staining. Where stated, 10 mM NH₄Cl (ammonium chloride; Thermo Fisher Scientific) was used. Cells were stained for 30 min at 4 °C with Live/Dead™ Near-IR dye (1:1000; Thermo Fisher Scientific), fixed with 2% paraformaldehyde for 10 min at room temperature, and permeabilized using Permeabilization Buffer (eBioscience) for 10 min at room temperature. Cells were then stained with CD3-PerCP-Cy5.5 (1:100; BioLegend), CD161-PE (1:200; Miltenyi), Vα7.2-PE-Cy7 (1:100; BioLegend), and vimentin-APC (1:50; Thermo Fisher Scientific).

Data acquisition was performed using a MACSQuant cytometer (Miltenyi) or an ImageStream system at the Dunn School of Pathology, University of Oxford (Cytek Biosciences). Data were analyzed using FlowJo software (Tree Star Inc.).

### Maintenance of cell lines

Cell lines were cultered at 37°C in 5% CO_2_. Caco2 cells (Colorectal adenocarcinoma cell line, ATCC), HHL12 cells (immortalized hepatocytes), TMNK1 (LSEC-like cells) were cultered at a starting density of 2×10^6^ cells in a T75 cell-culture flask, using GlutaMAX medium supplemented with 10% FBS, 1% MEM non-essential-amino acid solution (NEAA, Gibco™), 1% PS, 1% L-glutamine (Thermo Fisher Scientific). Cultures were kept monitored daily and split when confluency was around 70-80%.

MAIT cell lines were derived from healthy PBMCs isolated by density gradient centrifugation on Ficoll. The cells were FACS-sorted based on CD3 expression and MR1 Tetramer staining, then expanded through stimulation with phytohemagglutinin (purified PHA) and human IL-2 (100 IU/ml) in the presence of γ-irradiated PBMCs (35 Gray) as feeder cells. MAIT cells were maintained in RPMI 1640 medium supplemented with 5% AB+ human serum, IL-2 (100 IU/ml, PeproTech®), Penicillin/Streptomycin (100 IU/ml; P/S, ThermoFIsher), pyruvate (Thermo Fisher), and non-essential amino acids (NEAA, Gibco™), with restimulation occurring every 14 to 25 days. Cells used for experiments were at least 12 days post-restimulation.

### In-vitro wound healing assay

In-vitro wound healing assays were performed using a Wound Healing Assay Kit (Abcam). Caco2 and HHL12 cells were seeded at a density of 6×10^5^ cells/well in 24-well-plate and cultured with GlutaMAX medium supplemented with 10% FBS, 1% MEM non-essential-amino acid solution (NEAA), 1% PS, and 1% L-glutamine (Thermo Fisher Scientific), until 80-90% confluency. The cell suspension was added to the well with the plastic insert in place and after 24h (Caco2) or 13h (HHL12) inserts were removed and MAIT cell supernatants were added in a 1:4 dilution. Supernatants from both sorted MAIT and the MAIT cell line with or without anti-VEGFR-2 (ThermoFisher) and/or anti-VIM (Abcam) blocking antibodies were used in the wound-healing assays to functionally validate the tissue repair function and the importance of this specific signaling pathway. In control experiments, rVEGF-A (1ng/mL, 50ng/mL and 100ng/mL, ThermoFisher) or rVIM (100ng/mL and 250ng/mL, ThermoFisher) were added in the presence or without the blocking anti VEGFR-2 (0.6ug/mL, ThermoFisher) or neutralizing anti VIM (2.32ug/mL, Abcam). To observe the closure of the scratch, time-course imaging was carried out with pictures taken at different intervals for a total of 24h for Caco2 and 13h for HHL12.

A published, publicly available ImageJ plugin was used to detect wound edges and quantify the closure rate of the wound area^41^.

### Immunofluorescence imaging

20’000 endothelial cells (LSEC cell line – TMNK1) were seeded in a 4-well chamber. After 24h, supernatants from the MAIT cell line were added to LSEC in the presence and without anti-VEGFR2 or anti-VIM, and medium was used as a control. After 24h, immunofluorescence staining protocol was performed as follows: permeabilized with a mixed solution of acetone/methanol for 20 min at 4 degrees, blocked with 4% skim milk for 30 min at room temperature, and stained with Ki67-FITC (BD Bioscences) overnight. The day after 5ul of DAPI (Vector Laboratories) solution was added to each chamber, and pictures were acquired using the Olympus microscope BX53 and a 10x magnification. Staining with isotypes was performed as a negative control. Images were analyzed by ImageJ, and the Area of positive signal was considered.

### Analysis of supernatants

Supernatants were analysed for growth factor expression using a LEGENDplex™ Human Growth Factor Panel (BioLegend) and Human VEGF-A and VEGF-B ELISA kits (LSBio), following the manufacturers’ instructions. For vimentin detection, proteins were precipitated from supernatants by addition of 6.1 N trichloroacetic acid (1:4, TCA:supernatant), vortexed, and incubated at 4 °C for 30 min. Samples were centrifuged at 14,000 g for 10 min, supernatants were discarded, and pellets were washed up to twice with cold acetone. Pellets were briefly dried at 95 °C to remove residual acetone and resuspended in reducing sample buffer. Samples were denatured at 95 °C for 10 min, separated on 4-12% NuPAGE Bis-Tris gels at 180 V for 45 min, and transferred to PVDF membranes (Biorad). Membranes were blocked with 5% skimmed milk in TBS-T [(3% NaCl, 2% TRIS, 1mL Tween20 in PBS (Thermo Fisher Scientific)] and incubated overnight at 4 °C with anti-vimentin primary antibody (Millipore), followed by HRP-conjugated goat anti-mouse secondary antibody (Jackson ImmunoResearch). Protein bands were visualized using Amersham™ ECL Prime detection reagent (Cytiva) and analysed using Fiji ImageJ.

### Multiplex staining and imaging of human liver sections

Human liver tissue was collected and frozen in OCT under REC 21/YH/0206. Frozen tissue was sectioned and immediately fixed in ice-cold acetone for 5 minutes, followed by brief incubation in 95% and 70% ethanol (ice, 2-3 minutes) and washed in PBS 2 x 5 minutes. Sections were blocked with 10% normal goat serum (Thermo Fisher Scientific) in PBS for 1 hour at room temperature before staining with multiple rounds of fluorescently labelled antibodies. For multiplex imaging a CellScape Precise Spatial Multiplexing platform (Bruker Spatial Biology, 3350 Monte Villa Parkway, Bothell, WA 98021, USA) was used.

Antibody cocktails were diluted in PBS containing 1% normal goat serum. The first round of antibodies also contained1:100 diluted Human TruStain FcX (Fc receptor blocking solution, Biolegend).

Multiple antibody cocktails were run in iterative cycles with 30 minutes incubation at room temperature followed by washing cycles using PBS, imaging, beaching and re-staining. For the images shown in Figure 3 the following antibodies and dilutions were used:

| Antibody target | Fluorophore | Clone | Dilution | Company | Catalogue | Lot |
| --- | --- | --- | --- | --- | --- | --- |
| Vα7.2 | PE | 3C10 | 1:100 | Biolegend | 351706 | B307783 |
| VEGF | PE | REAL1004 | 1:250 | Miltenyi | 130-127-122 | 5240107468 |
| CD161 | PE | REA631 | 1:100 | Miltenyi | 130-113-596 | 5210606249 |
| Vimentin | PE | REA409 | 1:500 | Miltenyi | 130-123-997 | 5221205134 |

Nuclei were stained with Sytox Green nucleic acid stain (Invitrogen) diluted 1:100000 in PBS and incubated for 30 minutes.

Individual images were acquired using a Plan Apo 20X 0.80 NA lens with 0.8 mm FOV and stitched into a full image overview for each channel. Full images including stitched images for all channels were exported as OME.TIFF files. Images were analysed in QuPathv0.6. 040. To generate figures, the 32bit Cellscape ome.tiff file was opened in Fiji ImageJ2.16.0^41^ using the Bioformats Importer plugin^42^ virtual stacks as individual channels. Stacks were made from selected channels, cropped, merged and converted to RGB colour. Single channel greyscale images of individual cells were saved as single tiff files from Qupath.

## Acknowledgments

We thank Robert Hedley and Vasiliki Tsioligka for technical assistance with ImageStream analysis and support during the experiment at the Flow Cytometry Facility, Sir William Dunn School of Pathology, University of Oxford. The authors acknowledge Catherine De Lara and Claire Hutchings for their help with PBMC isolation and for scientific discussions related to this work. We also thank Matthew Edmands, Sam M. Murray, and Hossain Akther Delowar for valuable scientific discussions. 5-OP-RU was a gift from David Fairlie, University of Queensland. KS was supported by a Fondazione Cassa di Risparmio di Padova e Rovigo (CARIPARO) grant and by a European Federation of Immunological Societies and Immunology Letters (EFIS-IL) scholarship. PK was supported by Wellcome (222426/Z/21/Z), the NIH (U19 I082360), the NIHR Oxford Biomedical Research Centre and an NIHR Senior Fellowship. C.-P.H. was supported by a Medical Sciences Internal Fund (award 0009784.)

## Author contributions

Conceived and designed the experiments: K.S., M.L., K.P., L.C.G., A.B., C.-P.H., and P.K. Performed the experiments: K.S., M.L., K.P., I.T., A.B., N.R., C.-P.H., P.K. Analyzed the data: K.S., M.L., K.P., I.T., L.C.G., A.B., C.-P.H. Contributed to the writing of this manuscript: K.S., M.L., C.-P.H, P.K. Contributed to the reviewing and editing: K.S., M.L., K.P., I.T., L.C. G., A.B., N.R., F.P.R., C.-P.H., P.K.

## Data and Material availability

All data will be made available upon publication.

## References

1 Porcelli, S., Yockey, C. E., Brenner, M. B. & Balk, S. P. Analysis of T cell antigen receptor (TCR) expression by human peripheral blood CD4-8- alpha/beta T cells demonstrates preferential use of several V beta genes and an invariant TCR alpha chain. J Exp Med 178, 1–16 (1993). 10.1084/jem.178.1.1

2 Tilloy, F. et al. An invariant T cell receptor alpha chain defines a novel TAP-independent major histocompatibility complex class Ib-restricted alpha/beta T cell subpopulation in mammals. J Exp Med 189, 1907–1921 (1999). 10.1084/jem.189.12.1907

3 Treiner, E. et al. Selection of evolutionarily conserved mucosal-associated invariant T cells by MR1. Nature 422, 164–169 (2003). 10.1038/nature01433

4 Kurioka, A. et al. MAIT cells are licensed through granzyme exchange to kill bacterially sensitized targets. Mucosal Immunol 8, 429–440 (2015). 10.1038/mi.2014.81

5 Provine, N. M. & Klenerman, P. MAIT Cells in Health and Disease. Annu Rev Immunol 38, 203–228 (2020). 10.1146/annurev-immunol-080719-015428

6 Leng, T. et al. TCR and Inflammatory Signals Tune Human MAIT Cells to Exert Specific Tissue Repair and Effector Functions. Cell Rep 28, 3077–3091 e3075 (2019). 10.1016/j.celrep.2019.08.050

7 Constantinides, M. G. et al. MAIT cells are imprinted by the microbiota in early life and promote tissue repair. Science 366 (2019). 10.1126/science.aax6624

8 Hinks, T. S. C. et al. Activation and In Vivo Evolution of the MAIT Cell Transcriptome in Mice and Humans Reveals Tissue Repair Functionality. Cell Rep 28, 3249–3262 e3245 (2019). 10.1016/j.celrep.2019.07.039

9 du Halgouet, A. et al. Role of MR1-driven signals and amphiregulin on the recruitment and repair function of MAIT cells during skin wound healing. Immunity 56, 78–92 e76 (2023). 10.1016/j.immuni.2022.12.004

10 Lamichhane, R. et al. TCR- or Cytokine-Activated CD8(+) Mucosal-Associated Invariant T Cells Are Rapid Polyfunctional Effectors That Can Coordinate Immune Responses. Cell Rep 28, 3061–3076 e3065 (2019). 10.1016/j.celrep.2019.08.054

11 Linehan, J. L. et al. Non-classical Immunity Controls Microbiota Impact on Skin Immunity and Tissue Repair. Cell 172, 784–796 e718 (2018). 10.1016/j.cell.2017.12.033

12 Arpaia, N. et al. A Distinct Function of Regulatory T Cells in Tissue Protection. Cell 162, 1078–1089 (2015). 10.1016/j.cell.2015.08.021

13 Burzyn, D. et al. A special population of regulatory T cells potentiates muscle repair. Cell 155, 1282–1295 (2013). 10.1016/j.cell.2013.10.054

14 Walkenhorst, M. et al. Protective effect of TCR-mediated MAIT cell activation during experimental autoimmune encephalomyelitis. Nat Commun 15, 9287 (2024). 10.1038/s41467-024-53657-9

15 Kurioka, A., Walker, L. J., Klenerman, P. & Willberg, C. B. MAIT cells: new guardians of the liver. Clin Transl Immunology 5, e98 (2016). 10.1038/cti.2016.51

16 Lamichhane, R. et al. Human liver-derived MAIT cells differ from blood MAIT cells in their metabolism and response to TCR-independent activation. Eur J Immunol 51, 879–892 (2021). 10.1002/eji.202048830

17 Lett, M. J. et al. Stimulatory MAIT cell antigens reach the circulation and are efficiently metabolised and presented by human liver cells. Gut 71, 2526–2538 (2022). 10.1136/gutjnl-2021-324478

18 Li, Y. et al. Mucosal-Associated Invariant T Cells Improve Nonalcoholic Fatty Liver Disease Through Regulating Macrophage Polarization. Front Immunol 9, 1994 (2018). 10.3389/fimmu.2018.01994

19 Brazovskaja, A. et al. Cell atlas of the regenerating human liver after portal vein embolization. Nat Commun 15, 5827 (2024). 10.1038/s41467-024-49236-7

20 Lee, C. et al. Vascular endothelial growth factor signaling in health and disease: from molecular mechanisms to therapeutic perspectives. Signal Transduct Target Ther 10, 170 (2025). 10.1038/s41392-025-02249-0

21 Chen, R., Lee, C., Lin, X., Zhao, C. & Li, X. Novel function of VEGF-B as an antioxidant and therapeutic implications. Pharmacol Res 143, 33–39 (2019). 10.1016/j.phrs.2019.03.002

22 Bry, M., Kivela, R., Leppanen, V. M. & Alitalo, K. Vascular endothelial growth factor-B in physiology and disease. Physiol Rev 94, 779–794 (2014). 10.1152/physrev.00028.2013

23 Cudmore, M. J. et al. The role of heterodimerization between VEGFR-1 and VEGFR-2 in the regulation of endothelial cell homeostasis. Nat Commun 3, 972 (2012). 10.1038/ncomms1977

24 Parvanian, S., Coelho-Rato, L. S., Patteson, A. E. & Eriksson, J. E. Vimentin takes a hike - Emerging roles of extracellular vimentin in cancer and wound healing. Curr Opin Cell Biol 85, 102246 (2023). 10.1016/j.ceb.2023.102246

25 van Beijnum, J. R. et al. Extracellular vimentin mimics VEGF and is a target for anti-angiogenic immunotherapy. Nat Commun 13, 2842 (2022). 10.1038/s41467-022-30063-7

26 Eriksson, J. E. et al. Introducing intermediate filaments: from discovery to disease. J Clin Invest 119, 1763–1771 (2009). 10.1172/JCI38339

27 Garner, L. C. et al. Single-cell analysis of human MAIT cell transcriptional, functional and clonal diversity. Nat Immunol 24, 1565–1578 (2023). 10.1038/s41590-023-01575-1

28 Yanai, H. et al. Tissue repair genes: the TiRe database and its implication for skin wound healing. Oncotarget 7, 21145–21155 (2016). 10.18632/oncotarget.8501

29 Shigyo, M., Kuboyama, T., Sawai, Y., Tada-Umezaki, M. & Tohda, C. Extracellular vimentin interacts with insulin-like growth factor 1 receptor to promote axonal growth. Sci Rep 5, 12055 (2015). 10.1038/srep12055

30 Pall, T. et al. Soluble CD44 interacts with intermediate filament protein vimentin on endothelial cell surface. PLoS One 6, e29305 (2011). 10.1371/journal.pone.0029305

31 Satelli, A., Hu, J., Xia, X. & Li, S. Potential Function of Exogenous Vimentin on the Activation of Wnt Signaling Pathway in Cancer Cells. J Cancer 7, 1824–1832 (2016). 10.7150/jca.15622

32 LeCouter, J. et al. Angiogenesis-independent endothelial protection of liver: role of VEGFR-1. Science 299, 890–893 (2003). 10.1126/science.1079562

33 Wang, L. et al. Liver sinusoidal endothelial cell progenitor cells promote liver regeneration in rats. J Clin Invest 122, 1567–1573 (2012). 10.1172/JCI58789

34 Salou, M. & Lantz, O. A TCR-Dependent Tissue Repair Potential of MAIT Cells. Trends Immunol 40, 975–977 (2019). 10.1016/j.it.2019.09.001

35 Mehta, H., Lett, M. J., Klenerman, P. & Filipowicz Sinnreich, M. MAIT cells in liver inflammation and fibrosis. Semin Immunopathol 44, 429–444 (2022). 10.1007/s00281-022-00949-1

36 Wang, W. et al. Liver transplant-facilitated CD161(+)Valpha7.2(+) MAIT cell recovery demonstrates clinical benefits in hepatic failure patients. Nat Commun 16, 4022 (2025). 10.1038/s41467-025-59308-x

37 Bolte, F. J. et al. Intra-Hepatic Depletion of Mucosal-Associated Invariant T Cells in Hepatitis C Virus-Induced Liver Inflammation. Gastroenterology 153, 1392–1403 e1392 (2017). 10.1053/j.gastro.2017.07.043

38 Bottcher, K. et al. MAIT cells are chronically activated in patients with autoimmune liver disease and promote profibrogenic hepatic stellate cell activation. Hepatology 68, 172–186 (2018). 10.1002/hep.29782

39 Hegde, P. et al. Mucosal-associated invariant T cells are a profibrogenic immune cell population in the liver. Nat Commun 9, 2146 (2018). 10.1038/s41467-018-04450-y

40 Liu, Y. et al. Mucosal-Associated Invariant T Cell Dysregulation Correlates With Conjugated Bilirubin Level in Chronic HBV Infection. Hepatology 73, 1671–1687 (2021). 10.1002/hep.31602

41 Suarez-Arnedo, A. et al. An image J plugin for the high throughput image analysis of in vitro scratch wound healing assays. PLoS One 15, e0232565 (2020). 10.1371/journal.pone.0232565

