## Supplemental Figures for "MAIT cells derived ligands signal via VEGFR2 to promote tissue repair and liver regeneration"

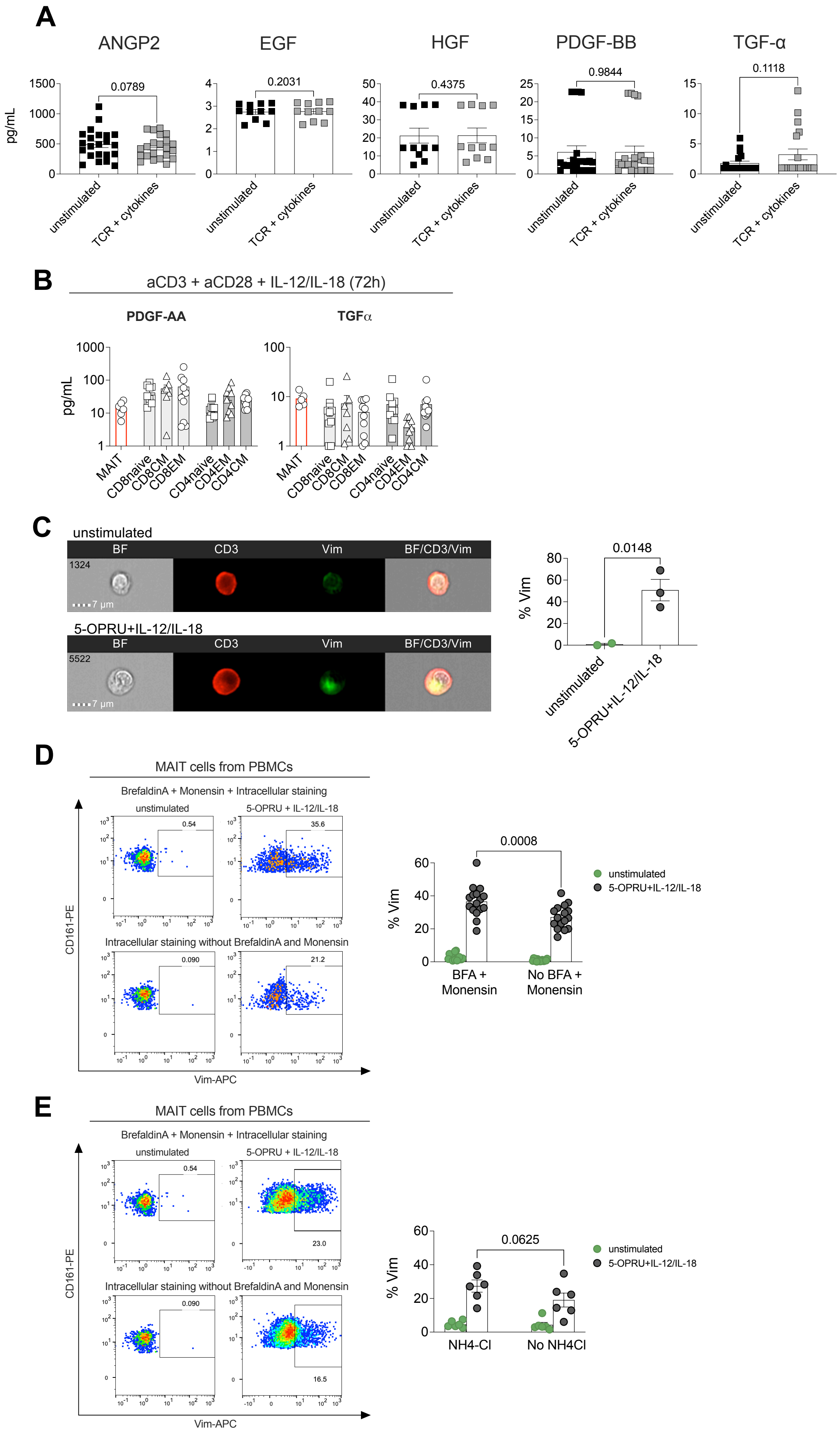
A
ANGP2
EGF
HGF
PDGF-BB
TGF-α
pg/mL
0.0789
0.2031
0.4375
0.9844
0.1118
unstimulated
TCR + cytokines
B
aCD3 + aCD28 + IL-12/IL-18 (72h)
PDGF-AA
TGFα
pg/mL
MAIT
CD8naive
CD8CM
CD8EM
CD4naive
CD4EM
CD4CM
C
unstimulated
BF
CD3
Vim
BF/CD3/Vim
1324
7 µm
5-OPRU+IL-12/IL-18
5522
0.0148
% Vim
D
MAIT cells from PBMCs
BrefaldinA + Monensin + Intracellular staining
5-OPRU + IL-12/IL-18
0.54
35.6
Intracellular staining without BrefaldinA and Monensin
0.090
21.2
CD161-PE
Vim-APC
0.0008
BFA + Monensin
No BFA + Monensin
5-OPRU+IL-12/IL-18
E
23.0
16.5
0.0625
NH4-Cl
No NH4Cl

**Supplementary Figure 1. MAIT cells secrete lower levels of additional growth factors compared with conventional T cells, and vimentin secretion is blocked by Brefeldin A and monensin but not by inhibitors of unconventional secretion pathways.**
(A) Concentration of the indicated growth factors secreted by primary blood-derived MAIT cells following activation with anti-CD3/anti-CD28 and IL-12/IL-18 for 72 hours, compared to unstimulated controls.
(B) Concentration of VEGF-A and GM-CSF in supernatants obtained from *in vitro* cultures of MAIT cells or the indicated conventional $CD8^+$ and $CD4^+$ T cell subsets after activation. Conventional T cells were classified as naïve ($CCR7^+$ $CD45RA^+$), central memory ($CCR7^+$ $CD45RA^-$), effector memory ($CCR7^-$ $CD45RA^-$), or terminally differentiated effector memory ($CCR7^-$ $CD45RA^+$).
(C) Peripheral blood mononuclear cells (PBMCs) were left unstimulated or stimulated for 72 hours with IL-12 (50 ng/mL) and IL-18 (50 ng/mL), 5-OP-RU (10 nM), or a combination of both stimuli. ImageStream flow cytometry was performed to assess vimentin expression in MAIT cells, defined as live $CD3^+$ $CD161^+$ $V\alpha7.2^+$ cells.
(D–E) PBMCs were left unstimulated or stimulated for 72 hours with IL-12 (50 ng/mL) and IL-18 (50 ng/mL), 5-OP-RU (10 nM), or a combination of both stimuli. Four hours prior to the end of stimulation, protein secretion was either left unblocked or inhibited using secretion inhibitors (Brefeldin A and monensin) or inhibitors of unconventional secretion pathways like $NH_4Cl$, followed by analysis of intracellular vimentin expression.

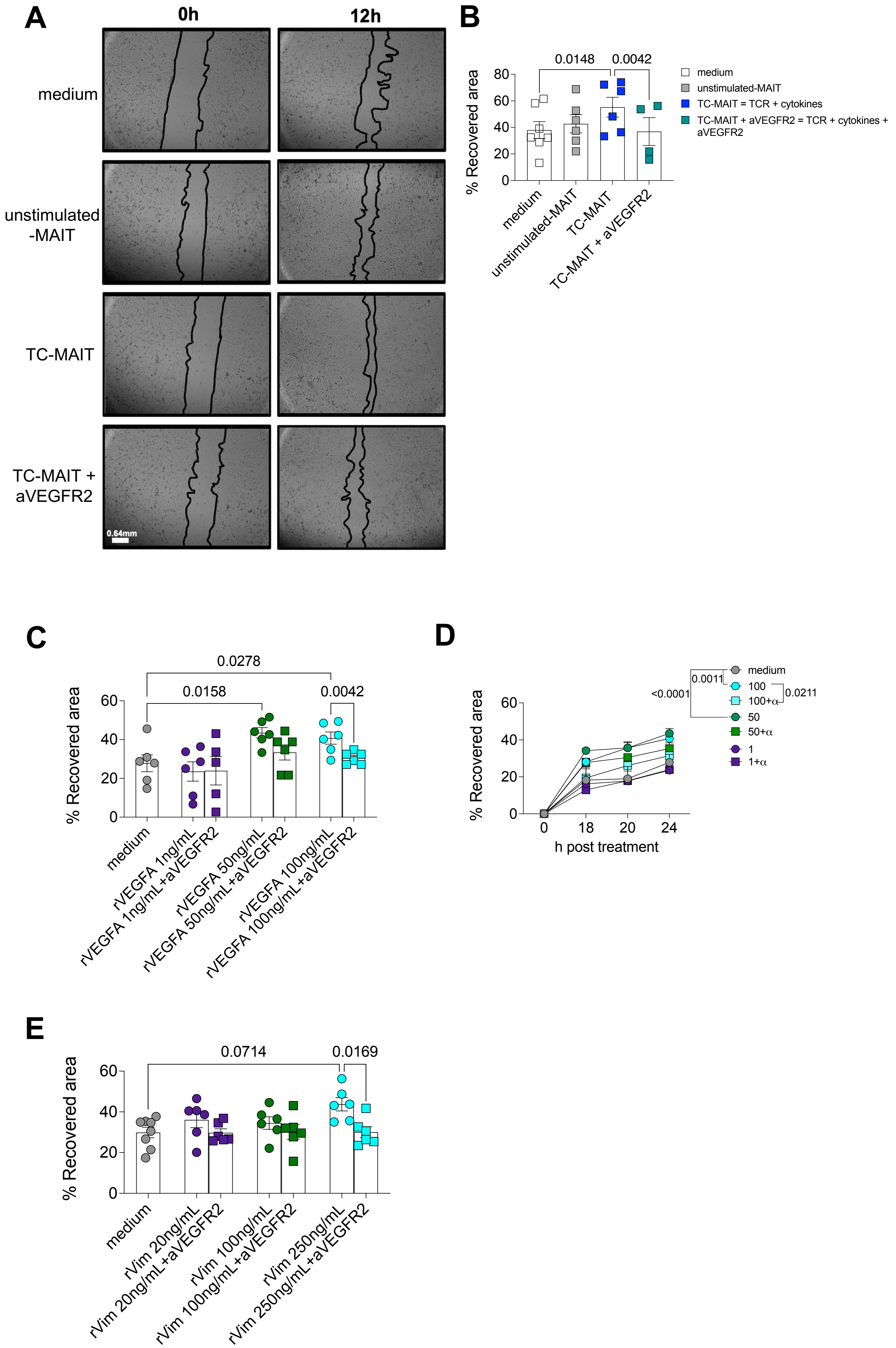
A
0h
12h
medium
unstimulated -MAIT
TC-MAIT
TC-MAIT + aVEGFR2
0.64mm
B
0.0148
0.0042
% Recovered area
medium
unstimulated-MAIT
TC-MAIT = TCR + cytokines
TC-MAIT + aVEGFR2 = TCR + cytokines + aVEGFR2
medium
unstimulated-MAIT
TC-MAIT
TC-MAIT + aVEGFR2
C
0.0278
0.0158
0.0042
% Recovered area
medium
rVEGFA 1ng/mL
rVEGFA 1ng/mL+aVEGFR2
rVEGFA 50ng/mL
rVEGFA 50ng/mL+aVEGFR2
rVEGFA 100ng/mL
rVEGFA 100ng/mL+aVEGFR2
D
0.0011
<0.0001
0.0211
medium
100
100+α
50
50+α
1
1+α
% Recovered area
h post treatment
E
0.0714
0.0169
% Recovered area
medium
rVim 20ng/mL
rVim 20ng/mL+aVEGFR2
rVim 100ng/mL
rVim 100ng/mL+aVEGFR2
rVim 250ng/mL
rVim 250ng/mL+aVEGFR2

**Supplementary Figure 2. RVEGF-A and rVIM promote accelaraton of wound closure , and effect is abrogated by anti-VEGFR2.**

(A, B) Quantification of wound closure, expressed as the percentage of recovered area, after 24 hours (A) or over the indicated time course (B) after treatment with different concentrations of rVEGFA with and without the presence of anti-VEGFR2.

(C) Quantification of wound closure, expressed as the percentage of recovered area, after 24 hours of using different concentrations of rVIM with and without the presence of anti-VEGFR2.

(D) Representative images of HHL12 cell monolayers before (0 h) and following treatment with MAIT cell–derived supernatants (12h) in the presence or absence of an anti-VEGFR2 antibody. Magnification is 10X.

(E) Quantification of wound closure, expressed as the percentage of recovered area, after 24 hours for each experimental condition.
